# Using a lentiviral Tet-regulated miR-E shRNA dual color vector to evaluate gene function in human leukemic stem cells in vivo

**DOI:** 10.1101/695965

**Authors:** Henny Maat, Jennifer Jaques, Edo Vellenga, Gerwin Huls, Vincent van den Boom, Jan Jacob Schuringa

**Affiliations:** Department of Experimental Hematology, Cancer Research Center Groningen, University Medical Center Groningen, University of Groningen, Hanzeplein 1, 9713 GZ Groningen, The Netherlands

## Abstract

RNA interference is a powerful tool to study loss-of-gene function in leukemic cells. Still, in order to identify effective novel treatment strategies to target and eradicate leukemic stem cells (LSCs), it is critically important to study gene function in a well-controlled and time-dependent manner. We implemented a lentiviral Tet-regulated miR-E shRNA dual color vector in our *in vitro* and *in vivo* human leukemia models. Thus, we were able to efficiently introduce doxycycline-inducible and reversible gene repression and trace and isolate transduced miR-E shRNA expressing cells over time. As proof of concept we focused on the non-canonical PRC1.1 Polycomb complex, which we previously identified to be essential for LSCs (1). Here, we show that inducible downmodulation of PCGF1 strongly impaired the growth of primary MLL-AF9 cells. Next, a Tet-regulated miR-E PCGF1 human xenograft MLL-AF9 leukemia mouse model was established, which revealed that early knockdown of PCGF1 at the onset of leukemia development significantly reduced peripheral blood chimerism levels and improved overall survival. In contrast, knockdown of PCGF1 when leukemia was already firmly established in the bone marrow proved insufficient to enhance overall survival. Despite these findings, FACS analysis of MLL-AF9/miR-E PCGF1/CD45^+^/GFP^+^ populations suggested that particularly cells with inefficient PCGF1 knockdown contributed to leukemogenesis. In conclusion, by building *in vivo* xenograft leukemia inducible RNAi models, we show that timing of gene knockdown critically impacts on the efficacy of leukemia treatment and that clonal drift still plays a major role in the escape of LSCs.

## Introduction

Leukemogenesis is a multistep process in which genetic and epigenetic changes affect normal growth and differentiation of hematopoietic stem and/or progenitor cells, which can ultimately lead to full leukemic transformation (2–5). With current therapies the majority of the blast population is readily eradicated, though the rare quiescent population of LSCs is more difficult to target and the major cause of relapse of the disease (6–8). In order to eradicate these LSCs, a better understanding of the molecular mechanisms underlying human leukemia development is needed.

Loss-of-function studies using RNA interference screens have shown to be very useful in identifying signaling pathways that are required for the proliferation and survival of leukemic cells (1, 9–11). Lentiviral short hairpin RNA (shRNA) expression vectors enable sustained and sequence-specific downregulation of a target mRNA (12). The usual setup of these RNA interference screens involves the transduction of leukemic cells with shRNAs resulting in quick downregulation of designated targets, after which phenotypes are evaluated *in vitro* or *in vivo* upon transplantation in (xenograft) mouse models. This however does not reflect the situation in the clinic at which leukemia patients are diagnosed and treated at the full-blown stage. Therefore, in order to evaluate the role of target genes in the maintenance and propagation of leukemic (stem) cells and to truly evaluate the efficacy of novel treatment options, one needs to first establish a leukemia within the bone marrow microenvironment and then perform gene-function studies.

The inducible Tet-regulated (Tet-On) shRNA expression system provides an attractive approach to study genes required for leukemic transformation (initiation) as well as for maintenance and survival (13–16). It allows controlled gene regulation by doxycycline inducible and reversible gene repression to study direct consequences at any chosen time point during disease progression. In addition, essential genes can be studied where constitutive knockdown might be toxic. To introduce Tet-regulated shRNA expression vectors in *in vivo* xenograft leukemia models would be very promising as a predictive model for therapy in (leukemia) patients. Treatment strategies and potential relapse of disease can be further studied upon discontinuing doxycycline treatment. Identifying and prospectively isolating different AML subclones expressing Tet-regulated shRNAs using leukemia specific plasma membrane markers can be used to study clone-specific dynamics in response to shRNA-mediated knockdown(de Boer et al, in submission).

In this study, we implemented a Tet-regulated miR-E shRNA expression vector, as reported by the Zuber lab, in our human *in vitro* and *in vivo* xenograft model systems to study gene function in the development and maintenance of leukemia (17). A single lentiviral dual-color vector allowed tracking of transduced miR-E shRNA expressing cells and demonstrated efficient inducible and reversible regulation of gene expression. Our data indicate that this doxycycline inducible knockdown approach is applicable to study the biological importance of a gene at any time in leukemia development *in vitro* and *in vivo.*

## Materials and Methods

### miR-E shRNA cloning

The lentiviral miR-E expression vector pRRL.TRE3G.GFP.miRE.PGK.mCherry.IRES.rtTA3 (LT3GECIR) was generously provided by the Zuber lab (17). miR-E shRNAs were designed for RING1B, PCGF1, KDM2B and a SCR control (see S1 Table) and generated as described by Fellmann et al (17). Briefly, 97-mer oligonucleotides were purchased from Invitrogen and PCR amplified using the primers miRE-Xho-fw (5’ TGAACTCGAGAAGGTATATTGCTGTTGACAGTGAGCG-3’) and miRE-EcoOligo-rev (5’-TCTCGAATTCTAGCCCCTTGAAGTCCGAGGCAGTAGGC-3’) according to Phusion High-Fidelity kit (Thermo Scientific). The PCR product (139 bp) was first ligated into pJet1.2 using blunt-end cloning and verified by sequencing. Then the miR-E shRNA was isolated using XhoI and EcoRI digestion and ligated into LT3GECIR vector also using XhoI and EcoRI digestion, resulting in the pRRL.TRE3G.GFP.miR-E shRNA.PGK.mCherry.IRES.rtTA3 construct.

### Lentiviral vectors and transductions

UMG LV6 MLL-AF9 mBlueberry2 was generated by swapping the AgeI-BsrGI mBlueberry2 fragment from the pLKO.1 mBlueberry2 vector into the UMG LV6 MLL-AF9 GFP vector (18), thereby replacing GFP with mBlueberry2. Generation of lentiviral viruses and transductions were performed as described previously (19). In addition, for primary cells miR-E shRNA viruses were concentrated to increase transduction efficiency using Centriprep Centrifugal filter YM-50 (Millipore, Billerica, MA, USA).

### Cell culture

K562, THP1 and NB4 cells (ATCC: CCL-243, TIB-202 and DSMZ: ACC 207) were maintained in RPMI 1640 (BioWhittaker, Lonza, Verviers, Belgium), 10% fetal bovine serum (FCS, HyClone Laboratories, Logan, Utah, US) and 1% penicillin/streptomycin (p/s, PAA Laboratories). Neonatal cord blood (CB) was obtained from healthy full-term pregnancies at the Obstetrics departments of Martini Hospital and University Medical Center Groningen (UMCG) after informed consent was obtained in accordance with the Declaration of Helsinki and approved by the UMCG Medical Ethical Committee. Mononuclear cells were isolated by density gradient centrifugation and CD34^+^ cells were subsequently isolated using the MACS CD34 MicroBead kit and the autoMACS (Miltenyi Biotec, Amsterdam, The Netherlands). CB-MLL-AF9 (here mBlueberry2) transformed cells were generated as described before (1, 20) and cultured in Gartner’s medium supplemented with 20 ng/ml Flt-3L, stem cell factor (SCF) and IL-3 in liquid conditions. Gartner’s medium consisted of alpha-MEM (BioWhittaker) supplemented with 12.5% heat-inactivated FCS and 12.5% heat-inactivated Horse serum (Sigma-Aldrich, Saint Louis, MO, USA), 1 μM hydrocortisone (Sigma-Aldrich), 57.2 μM β-mercaptoethanol (Sigma) and 1% p/s. Doxycycline was added in a concentration of 1 μg/ml every 2-3 days.

### RNA isolation and quantitative real-time PCR

Total RNA was isolated using the RNeasy Mini Kit from Qiagen (Venlo, The Netherlands). For quantitative RT-PCR, RNA was reverse transcribed using the iScript cDNA synthesis kit (Bio-Rad) and amplified using SsoAdvanced SYBR Green Supermix (Bio-Rad) on a CFX384 Touch Real-Time PCR Detection System (Bio-Rad). RPL27 was used as housekeeping gene. Primer sequences are listed in S2 Table.

### Flow cytometry analysis

Flow cytometry analyses were performed on the BD LSR II (Becton Dickinson (BD) Biosciences) and data were analyzed using FlowJo (Tree Star Inc, Ashland, OR, USA). For peripheral blood, bone marrow (BM), spleen (SP) and liver (LV) analysis, stains for CD45-APC (HI30) and CD19-BV785 (HIB19) (Biolegend) were included. Prior to staining, cells were blocked with anti-human FcR block (MACS miltenyi Biotec) and anti-mouse CD16/CD32 block (BD Biosciences). For cell sorting the MoFlo XDP or MoFLo Astrios (Beckman Coulter) were used.

### Inducible miR-E shRNA expression in a xenograft MLL-AF9 model

Eight to ten week old female NSG (NOD.Cg-Prkdcscid ll2rgtm1Wjl/SzJ) mice were purchased from the Centrale Dienst Proefdieren (CDP) breeding facility within the University Medical Center Groningen. Mouse experiments were performed in accordance with national and institutional guidelines, and all experiments were approved by the Institutional Animal Care and Use Committee of the University of Groningen (IACUC-RuG). Mice were sub-lethally irradiated with a dose of 1 Gy (X-RAD 320 Unit, PXINC 2010) 24 hours before IV (lateral tail vein) injections. For secondary transplantations, mice were injected with 5.10^4^ MLL-AF9 (mBlueberry)/miR-E PCGF1 (mCherry^+^) cells. After irradiation mice received Neomycin (3.5 g/l) in their drinking water for two weeks. Doxycycline was administered by feeding mice doxycycline-containing food pellets (625ppm, Special Diet Services, Witham, England) and replenished every week. Peripheral blood chimerism levels were monitored by regular blood sample analysis. Mice were terminated by cervical dislocation under isoflurane anesthesia when chimerism levels in the blood exceeded 30% (MLL-AF9/miR-E PCGF1/CD45^+^). Peripheral blood, BM, SP and LV were analyzed by FACS.

## Results

### An efficient lentiviral Tet-regulated miR-E shRNA expression vector in stably transduced leukemic cells

In order to study the function of genes necessary for the maintenance and survival of leukemic cells *in vitro* and *in vivo* in our human models, we took advantage of a Tet-regulated (inducible) lentiviral miR-E expression vector reported by the Zuber lab (17). We generated miR-E based shRNAs against PRC1.1 subunits RING1B, PCGF1 and KDM2B, plus a SCR control, that were cloned into the pRRL.TRE3G.GFP.miRE.PGK.mCherry.IRES.rtTA3 vector (LT3GECIR) (Fig 1A). These targets were selected based on our previous studies in which we identified non-canonical PRC1.1 as a critically important Polycomb complex in human leukemic cells (1). In K562 cells, the transduction efficiency reached ~40% and mCherry^+^ cells were sorted to establish stable cell lines. The tetracycline-derivate doxycycline (dox, 1 μg/ml) was used to induce GFP-coupled miR-E shRNA expression and therefore could be easily traced. Dox-induced shRNA expression was very efficient as demonstrated by >95% mCherry^+^/GFP^+^ cells (Fig 1A). To test the efficiency of the knockdown, qRT-PCR analysis was performed on K562 cells expressing miR-E RING1B, miR-E PCGF1 or miR-E KDM2B in the presence or absence of doxycycline for 4 days (Fig 1B). Next, we examined the effect of dox withdrawal to verify that the knockdown was reversible (Fig 1C). In the presence of dox, miR-E KDM2B cells became GFP-positive within two days, coinciding with a 60% knockdown in KDM2B expression (Fig 1C). In order to evaluate the reversibility of the system, cells were withdrawn from dox. Already after two days of dox withdrawal the GFP percentage started to drop, but it took 10 days before all GFP expression was completely gone. At that timepoint, KDM2B expression levels were also again up to normal levels (Fig 1C). Thus, these lentiviral vectors provide an efficient dox-inducible and reversible shRNA expression tool, which is useful to perform gene function studies in a timedependent manner.

**Fig 1.**
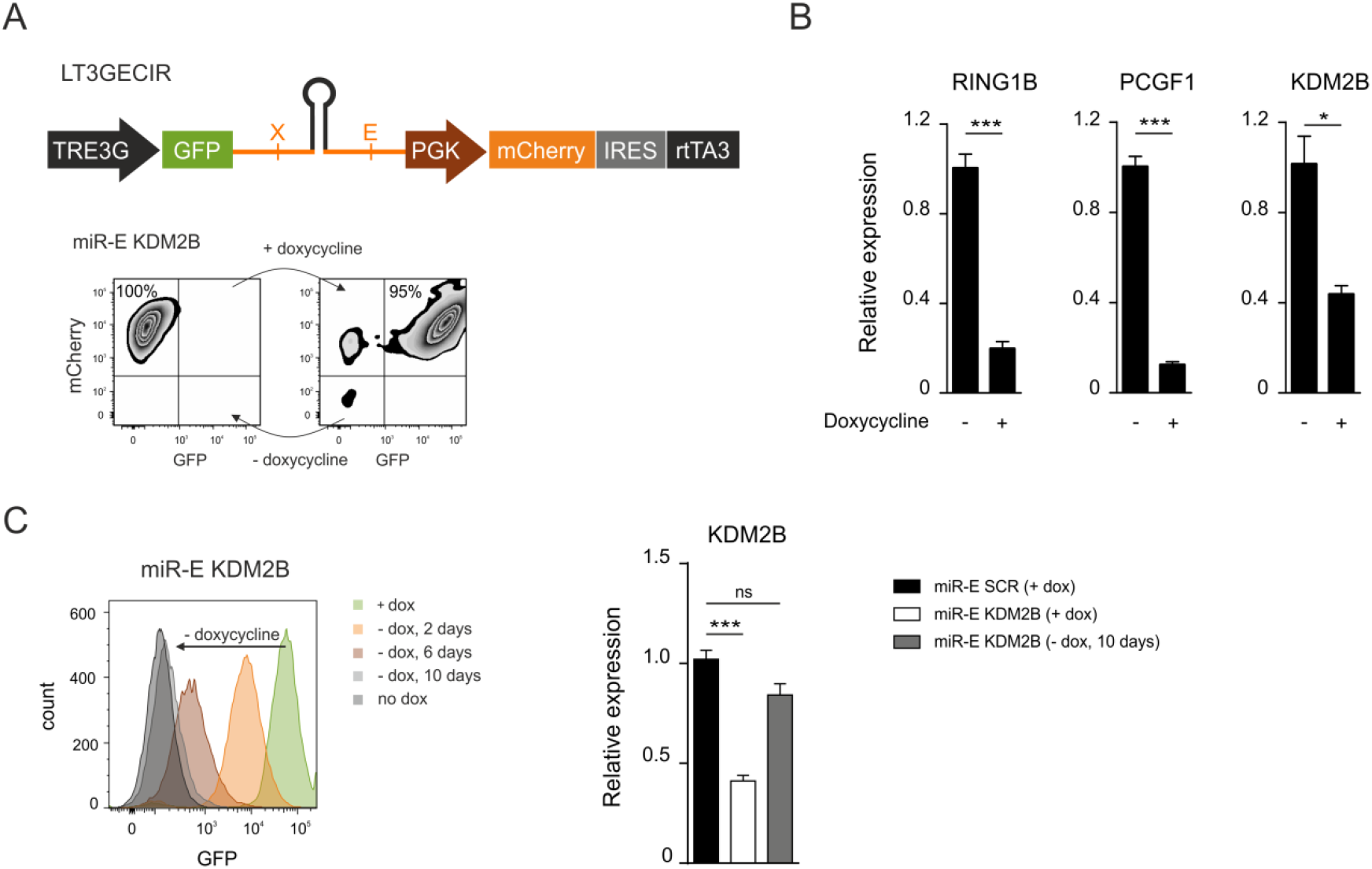
Inducible and reversible miR-E shRNA expression *in vitro.* (A) Schematic representation of the lentiviral Tet-regulated miR-E shRNA expression vector (illustration adapted from Fellmann et al., 2013) expressing fluorescence (mCherry/GFP) coupled miR-E shRNAs. As a representative example, doxycycline induced shRNA expression efficiency (*in vitro)* is shown as analyzed by FACS in miR-E KDM2B min/plus dox. (B) Knockdown efficiency of RING1B, PCGF1 and KDM2B in K562 miR-E expressing shRNAs respectively, with and without doxycycline for 4 days. Error bars represent SD. (C) Flow cytometry analysis of K562 miR-E KDM2B cells in the presence and/or withdrawal of doxycycline. The mean fluorescence intensity (MFI) for GFP, as a marker for doxycycline induced shRNA expression, is plotted for different time points. KDM2B expression was analyzed by qRT-PCR. Statistical analysis was performed using an unpaired t-test (p≤ 0.05 (*), p ≤ 0.01 (**) and p≤0.001 (***).

### Primary human MLL-AF9 cells are sensitive to inducible downmodulation of PRC1.1 components

Next, we examined the biological importance of downregulating PRC1.1 subunits RING1B, PCGF1 and KDM2B in human leukemic cells *in vitro.* First, AML cell lines NB4 (PML-RARA) and THP-1 (MLL-AF9) were transduced with the miR-E KDM2B shRNA vector. Upon dox treatment the majority of mCherry^+^ cells became GFP^+^, indicating efficient miR-E induction (S1 Fig). Following three weeks, we observed a clear reduction in mCherry^+^/GFP^+^ cells in the miR-E KDM2B transduced group indicating that cells rely on KDM2B function (S1 Fig). Furthermore, we setup a primary cord blood (CB)-MLL-AF9 model expressing miR-E PRC1.1 inducible shRNAs (Fig 2A). CB derived CD34^+^ cells were transduced with the lentiviral MLL-AF9 mBlueberry vector. Then, transformed MLL-AF9 cells were transduced with lentiviral constructs for miR-E SCR, miR-E RING1B, miR-E PCGF1 and miR-E KDM2B, reaching transduction efficiencies of 10-15% (Fig 2B). While dox-induced mCherry^+^/GFP^+^ percentages remained stable over time in the miR-E SCR transduced control group, downregulation of either RING1B, PCGF1 or KDM2B resulted in impaired growth of primary MLL-AF9 cells (Fig 2C and 2D).

**Fig 2:**
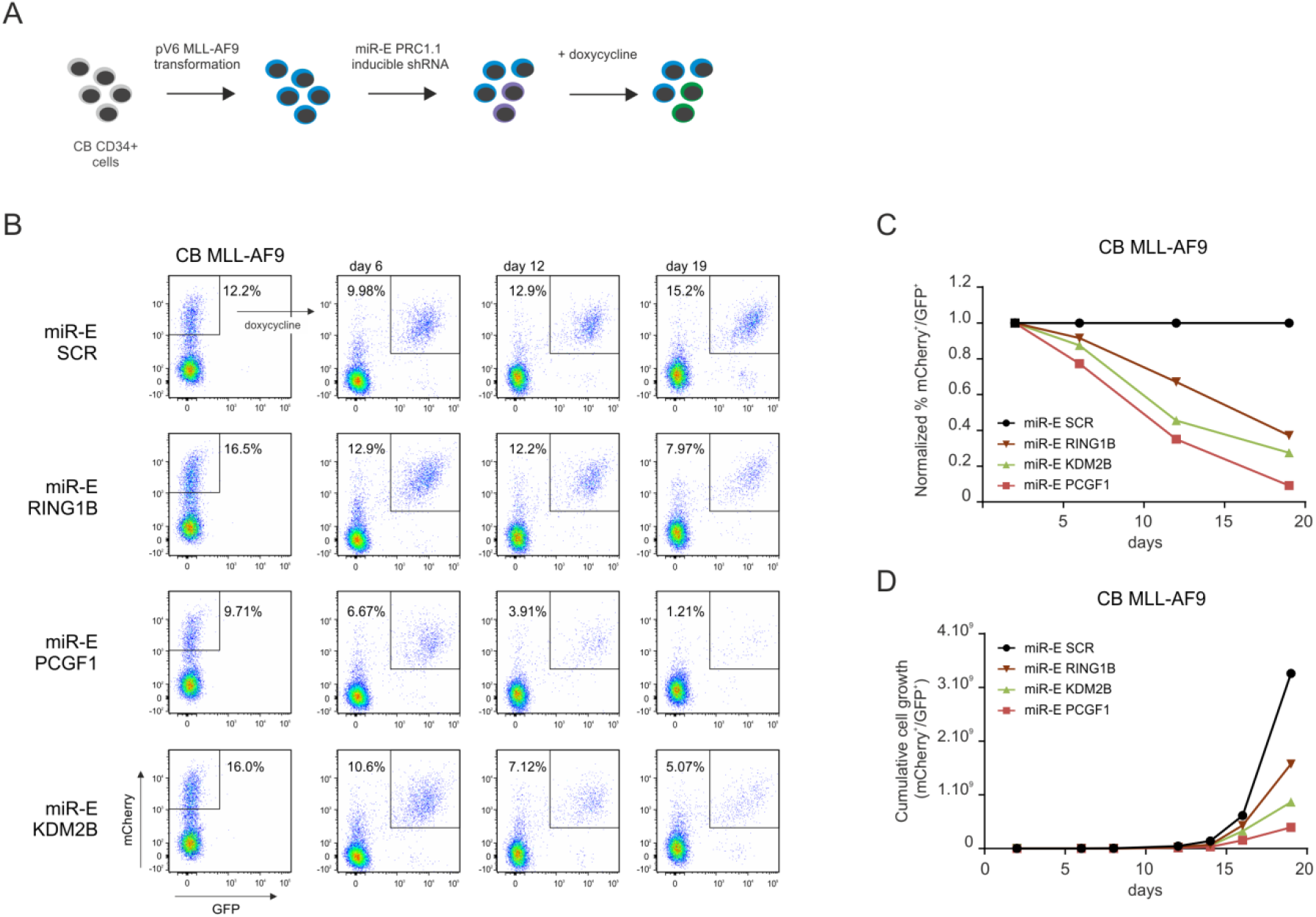
Doxycycline induced miR-E RING1B, PCGF1 or KDM2B shRNA expression impaired cell growth of primary human MLL-AF9 cells. (A) Experimental setup of primary CB MLL-AF9 (mBlueberry) miR-E shRNA (SCR, RING1B, PCGF1 and KDM2B) transductions. (B) Flow cytometry analysis of primary CB MLL-AF9 miR-E SCR, RING1B, PCGF1 and KDM2B cells in the presence of doxycycline over the course of 19 days. (C) Normalized percentage mCherry^+^/GFP^+^ and (D) cumulative cell growth (mCherry^+^/GFP^+^) of CB-MLL-AF9 cells expressing miR-E SCR, RING1B, PCGF1 or KDM2B.

### Evaluation of Tet-regulated miR-E shRNA expression in a xenograft MLL-AF9 model

To examine loss-of-function in a time-dependent manner *in vivo*, we developed a xenograft MLL-AF9 Tet-regulated miR-E shRNA mouse model. Since PCGF1 knockdown strongly reduced the proliferation of MLL-AF9 cells *in vitro*, we studied the consequences of PCGF1 knockdown at several stages during the process of leukemia development *in vivo*. CB-derived CD34^+^ cells were transduced with the lentiviral vector expressing miR-E PCGF1 and mCherry^+^ cells (~17% efficiency) were sorted. Subsequently, cells were transduced with the lentiviral MLL-AF9 (mBlueberry) vector and were intravenously injected into sub-lethally irradiated NSG mice (Fig 3A). After 30 weeks, leukemia developed with high levels of engraftment of MLL-AF9/miR-E PCGF1/CD45^+^/CD19^+^ cells in BM, SP and LV (S2A Fig). Next, secondary transplantations were performed and to knockdown PCGF1, doxycycline was administered to the mice in food pellets (625 ppm) either at the start or at 10 weeks after secondary transplantation when leukemia was already established in the BM (Fig 3B). Control mice that did not receive any doxycycline, developed leukemia within 150 days (Fig 3C). Mice that were treated with doxycycline from the start showed significant prolonged survival until the end of the experiment (Fig 3C). Peripheral blood chimerism levels (MLL-AF9/miR-E PCGF1/CD45^+^) remained low over time (Fig 3D). To investigate if we could treat mice at a later stage in the development of leukemia, we also started treatment of doxycycline 10 weeks after secondary transplantation when average blood chimerism levels were already 5%, a situation that would more faithfully mimic the patient situation (Fig 3E). While chimerism levels increased over time in untreated mice after week 10, this increase was clearly delayed in mice treated with doxycycline (Fig 3E). Nevertheless, leukemia did develop in mice treated from week 10 and mice were sacrificed when peripheral blood chimerism levels reached 30% and ultimately within this experiment no significant differences in overall survival were reached (Fig 3C, 3D and 3E).

**Fig 3.**
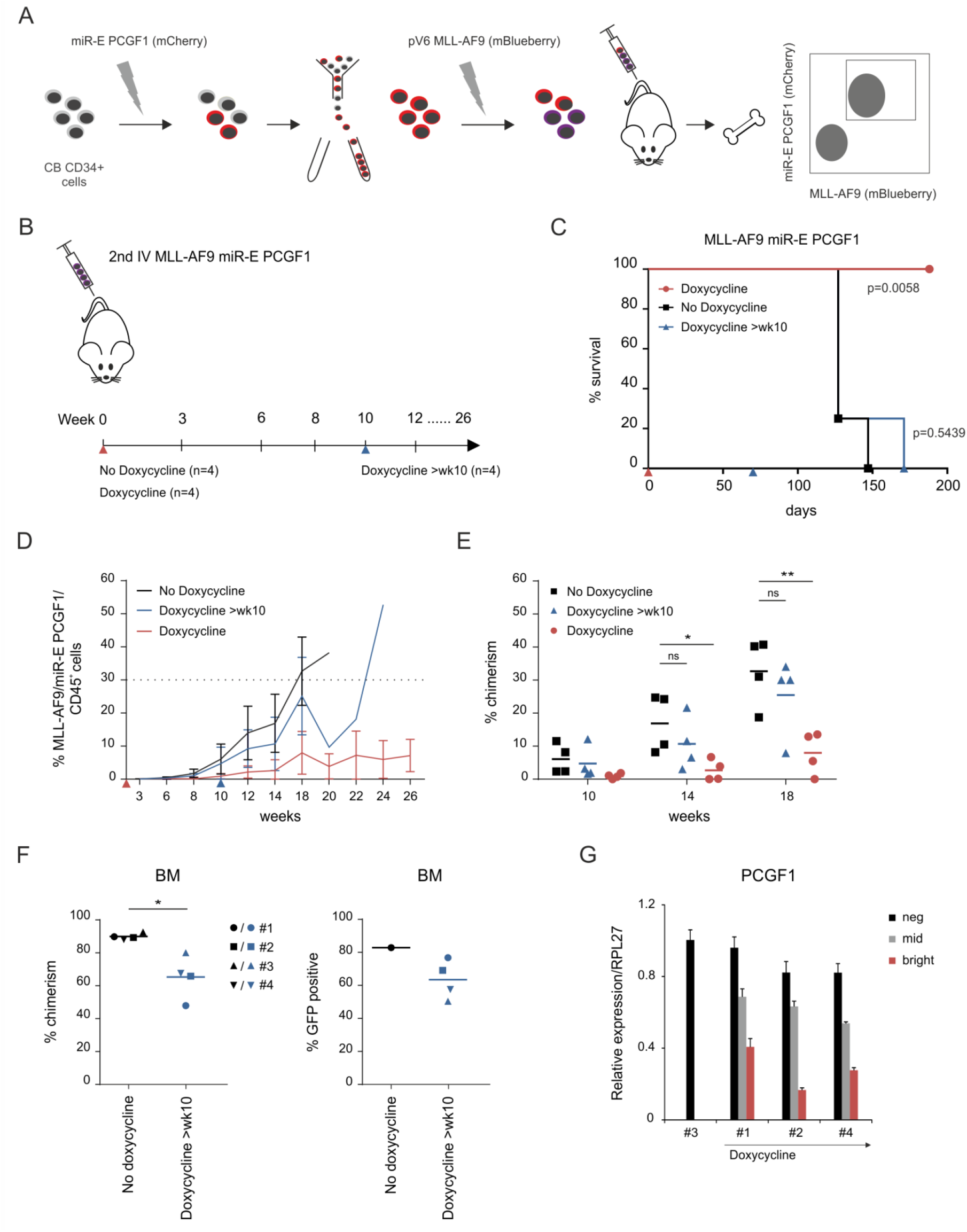
Evaluation of miR-E PCGF1 in xenograft CB-MLL-AF9 mouse model. (A) Schematic overview of developing an inducible miR-E shRNA against PCGF1 in a primary CB-MLL-AF9 (mBlueberry) *in vivo* model. (B) Experimental setup as illustrated where MLL-AF9 miR-E PCGF1 secondary transplantations were performed. Blood chimerism levels were monitored for indicated time points (weeks). Mice were divided into three groups, no doxycycline (n=4), doxycycline from the start (n=4) and doxycycline >wk10 (n=4). Doxycycline was administered in food pellets (625 ppm). (C) Kaplan-Meier curve of MLL-AF9 miR-E PCGF1 mice treated with no doxycycline, doxycycline from the start or doxycycline >wk10. Statistical analysis was performed using a log-rank test. (D) Peripheral blood chimerism levels plotted over the course of the experiment between no doxycycline, doxycycline from the start (red triangle) and doxycycline >wk10 (blue triangle). Mice were sacrificed when chimerism levels exceeded 30%. (E) Peripheral blood chimerism levels of individual mice treated from the start or treated for 4-8 weeks. Statistical analysis was performed using unpaired t-test (p ≤ 0.05 (*), p ≤ 0.01 (**)). (F) Engraftment levels in BM shown as % chimerism of MLL-AF9 mBlueberry/miR-E PCGF1 mCherry/ CD45^+^ cells in no doxycycline and doxycycline>wk10 treated mice. Within the population of engrafted cells, GFP^+^ cells are shown as a percentage. From the no doxycycline group, mouse #1 was administered doxycycline for 5 days before sacrifice to determine max GFP induction in the BM. (G) Expression of PCGF1 in BM samples of no doxycycline and doxycycline treated mice sorted on neg, mid and bright GFP populations, analyzed by qPCR.

Even though substantial levels of ~65% chimerism (MLL-AF9/miR-E PCGF1/CD45^+^) were observed in the BM of mice treated with doxycycline at week 10, these levels were still significantly lower compared to control mice (Fig 3F). Intriguingly, FACS analysis of MLL-AF9/miR-E PCGF1 mCherry^+^/CD45^+^ populations in doxycycline-treated mice revealed strong variation in GFP positivity (Fig 3F and S2B Fig). We questioned whether *in vivo* all BM cells would be equally sensitive to dox treatment as compared to *in vitro* conditions. To do so, a mouse (#1) that had not been treated with dox and had developed leukemia was administered with dox for 5 days before sacrifice. 82% of these cells were GFP positive and while this was somewhat lower compared to the 95% that was reached *in vitro*, this was still higher than percentages seen in mice that were treated with doxycycline for 8-14 weeks, which showed average GFP levels of 63% (Fig 3F). These data suggest that there was some selection for those clones that did not express GFP, or only at very low levels. Indeed, the highest chimerism levels were seen in mice with the lowest percentage of GFP^+^ cells. Moreover, we sorted the bright, mid and negative GFP populations and analyzed the efficiency of PCGF1 knockdown by qRT-PCR (Fig 3G). The GFP negative population showed similar PCGF1 expression levels as the no doxycycline control group, PCGF1 knockdown was more efficient in the GFP^mid^ population and the highest knockdown was achieved in the GFP^bright^ population (Fig 3G). Thus, these data indeed indicate that it were particularly those cells with inefficient or no PCGF1 knockdown that contributed to MLL-AF9-induced leukemogenesis. Re-analysis of the BM sample of the control mouse (#1) that was administered with dox for 5 days revealed that the highest miR-E PCGF1 mCherry^+^ cells (top) all expressed GFP, whereas the middle and bottom mCherry^+^ population represented GFP levels of 84% and 60% respectively (S2C Fig). These data suggest that the maximal dox-induced miR-E shRNA expression was technically limited to the high mCherry^+^ cells. Thus, in future studies to establish xenograft leukemia models expressing a dox-inducible miR-E shRNA vector, it is advisable to select the highest mCherry+ cells to reach maximal gene knockdown efficiencies.

## Discussion

Acute myeloid leukemia (AML) is characterised by (sub)clonal expansion of immature blasts in the bone marrow with a block in differentiation (21, 22). Whole blood cell counts, cytogenetics, flow cytometry and immunohistochemistry tests are instrumental in the diagnosis and classification of AML subtypes and subsequently treatment strategy (23, 24). Nevertheless, with currently available therapies the majority of patients will eventually still relapse. This is most likely due to the existence of therapy-resistant LSCs and many studies now focus on ways to identify, target and ultimately eradicate these LSCs in order to reach more curative therapies (25–29). To understand the molecular mechanisms involved in initiation, maintenance and relapse of human leukemias, it is essential to study gene function in a time-dependent manner in *in vitro* and *in vivo* human leukemic models (30, 31). Loss-of-gene function studies using constitutive RNAi expression result in a stable and quick downregulation of a designated target and while this is a powerful tool it will not reveal gene functions at specific stages of the disease. Therefore, an inducible RNAi system in a human leukemia model would allow for a controlled regulation of gene expression and be more promising as predictive model for therapy in patients. Where genetic knockout of essential genes using CRISPR-Cas9 might be too toxic for the cells, we reasoned that the best model would be an inducible knockdown system. While the application of CRISPR-Cas9 is very interesting, it remains a major challenge in primary AMLs (32, 33). We implemented an inducible lentiviral miR-E shRNA dual-color vector in our *in vitro* and *in vivo* human leukemia models that can be used to study the function of genes during any stage of leukemia development. This single lentiviral vector (LT3GECIR) based on an optimized miR-E backbone was shown to enhance shRNA levels and improve knockdown efficiency (17, 34). The dualcolor vector allows identification of mCherry^+^ transduced cells and in the presence of doxycycline, GFP-coupled miR-E shRNA is expressed. We showed that doxycycline-inducible miR-E shRNA expression in leukemic cells was very efficient *in vitro*, resulting in effective gene repression and the effects were also reversible. Next, we demonstrated that the effects of a dox-inducible lentiviral miR-E shRNA on cell proliferation can be studied in AML cell lines and primary MLL-AF9 cells. Downregulation of Polycomb proteins RING1B, PCGF1 or KDM2B resulted in impaired growth of primary MLL-AF9 cells indicated by a strong reduction in mCherry^+^/GFP^+^ cells, which is in line with our previous constitutive shRNA reported data in which PRC1.1 was found to be essential for leukemic cells (1). Furthermore, we introduced the miR-E shRNA vector in a xenograft MLL-AF9 mouse model. Doxycycline was administered in food pellets because it is a reliable and effective method (35, 36). In the BM, GFP-coupled miR-E shRNA levels reached maximum levels of 82%, indicating that the system is also nicely dox-inducible *in vivo,* albeit to a somewhat lesser extent compared to *in vitro* experiments where >95% GFP-positivity was reached. In mice carrying MLL-AF9 miR-E PCGF1 cells that were treated from the start with dox directly after transplantation, a significantly prolonged survival was observed. Chimerism levels in the blood remained low over time, indicating that the model can be used to study gene function *in vivo* and also underlining the importance of non-canonical PRC1.1 in leukemia development.

In contrast, and somewhat unexpectedly, the survival of mice in which dox treatment was initiated at week 10 was not significantly prolonged. We decided to start the treatment of mice when peripheral blood chimerism levels had reached about 5%, which in our view would be a moment that would faithfully recapitulate the situation when patients would enter the clinic. These data strongly contrast our *in vitro* data in which we observed that PCGF1 knockdown strongly impaired MLL-AF9 cell growth, and also the effects seen *in vivo* when mice were treated directly after transplantation, which were very promising. However, once leukemia was established in the BM niche the effects were minor, which would have major implications for the efficacy of PCGF1 inhibition as a treatment modality. We also observed that doxycycline treated mice with low chimerism levels in the blood, still had substantial levels of engraftment in the BM, suggesting that the niche would provide protective factors that counteract loss of PCGF1. Clearly, the bone marrow microenvironment might play a protective role in the survival of leukemic cells (37–39). Indeed, we observed that MLL-AF9 cells cultured in the presence of murine MS5 stromal cells were less sensitive towards USP7 inhibition compared to cells grown in liquid culture conditions (Maat et al, in submission). This even further emphasises that loss-of-gene function or inhibitor studies should be investigated in the context of a niche.

Intriguingly, we noted strong variations in GFP positivity in FACS analysis of MLL-AF9/miR-E PCGF1 mCherry^+^/CD45^+^ populations in doxycycline-treated mice. Therefore, we questioned whether in this *in vivo* context all BM cells would be equally sensitive to dox treatment compared to *in vitro* conditions. We indeed noted that, *in vivo*, the dox-inducibility was somewhat less compared to what was seen *in vitro*. Perhaps more importantly, we also noted that knockdown was most efficient in the brightest mCherry^+^/GFP^+^ cells, and the lower 40% of the mCherry^+^ fraction did not express GFP at all and evaded shRNA expression. Also in the THP1 cell line there was a minor mCherry^+^ population that did not become GFP^+^ upon doxycycline treatment. This indicated that certain cells were insensitive for doxycycline potentially due to failure to undergo proper tetracycline-responsive element expression, as also experienced by Zuber and colleagues (40). As a consequence, clonal selection had occurred in our mice treated with dox from week 10, whereby a shift towards GFP^mid/neg^ cells was seen with considerable less efficient knockdown of PCGF1. Thus, in future studies to establish xenograft leukemia models expressing a dox-inducible miR-E shRNA vector, it is advisable to select the highest mCherry^+^ cells to reach maximal gene knockdown efficiencies. With this in mind, we conclude that the inducible miR-E shRNA dual-color vector can be used to study, trace and isolate shRNA expressing cells in a time-dependent manner both in *in vitro* and *in vivo* human leukemia models that most faithfully recapitulate human disease.

## Supporting information

miR-E shRNAs

primer sequences

## Acknowledgments

The authors thank Dr. Johannes Zuber (Research Institue of Molecular Pathology (IMP), Medical University of Vienna, Vienna, Austria) for providing the pRRL.TRE3G.GFP.miRE.PGK.mCherry.IRES. rtTA3 (LT3GECIR) construct. We thank Bart-Jan Wierenga for help with cloning. We also thank Roelof-Jan van der Lei, Henk Moes, Theo Bijma en Geert Mesander for help with cell sorting. We greatly appreciate the help of Dr. J.J. Erich and Dr. A. van Loon and colleagues (Departments of Obstetrics, UMCG and Martini Hospital) for collecting CB. This work is supported by the European Research Council (ERC-2011-StG 281474-huLSCtargeting).

## Supporting Information

**S1 Table. miR-E shRNAs, 97-mer oligonucleotides**

**S2 Table. Primer sequences**

**S1 Fig.**
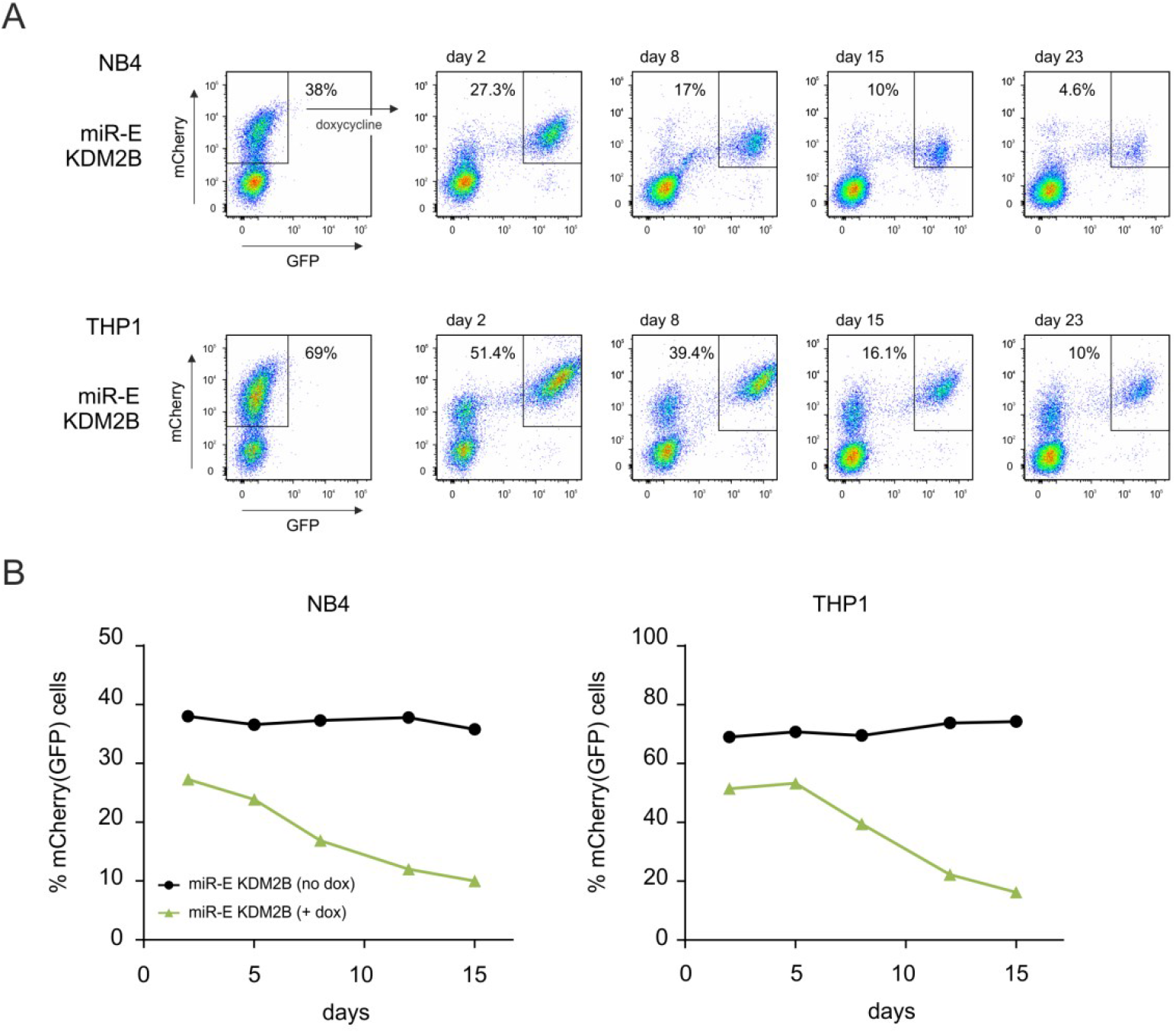
Inducible KDM2B knockdown impairs cell growth of leukemic cells. (A) Flow cytometry analysis of NB4 and THP1 miR-E KDM2B cells, unsorted, following doxycycline treatment for 3 weeks. (B) Analysis of miR-E KDM2B mCherry^+^ (no dox) or mCherry^+^/GFP^+^ NB4 or THP1 cells in time.

**S2 Fig.**
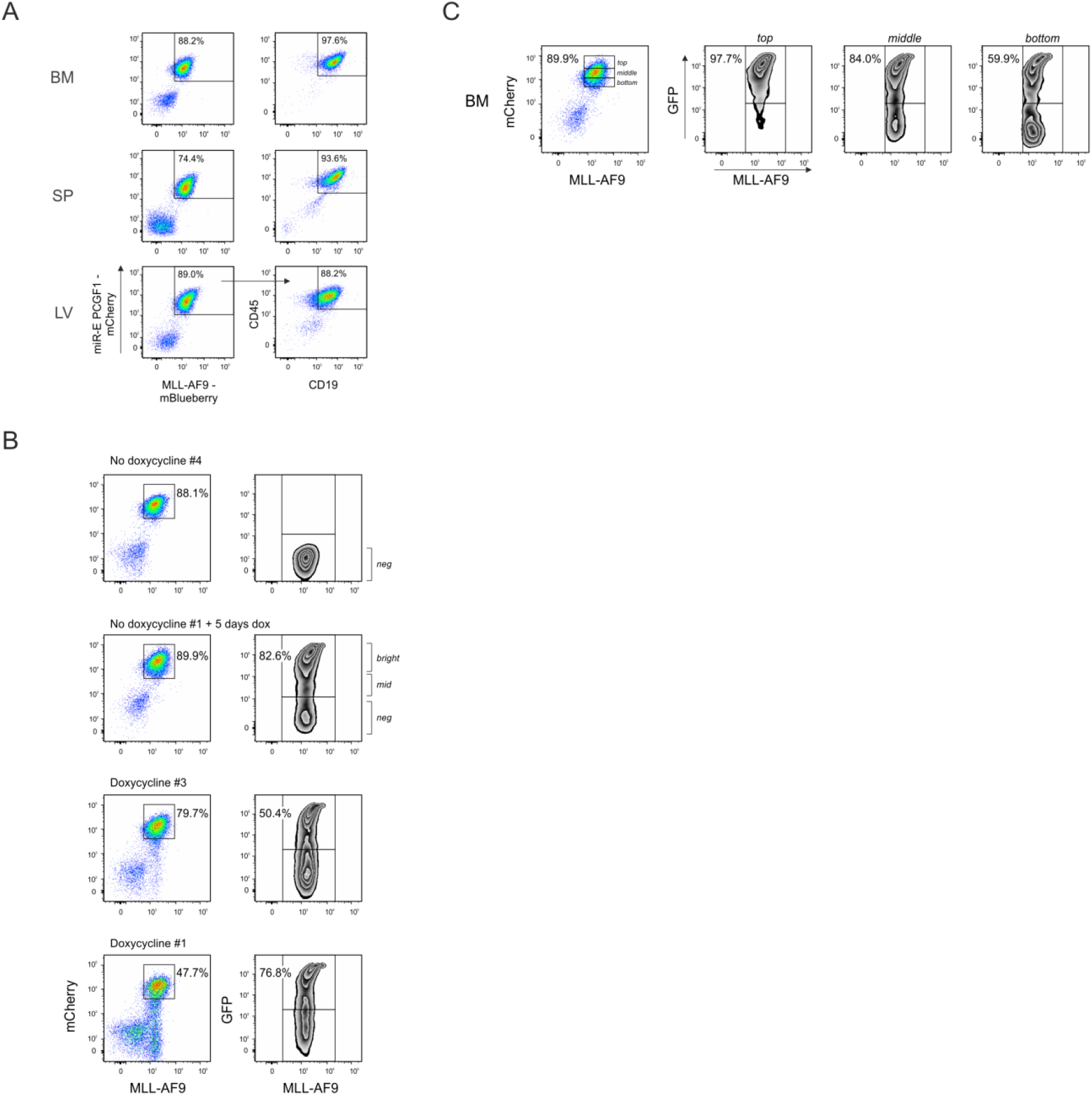
FACS analysis of MLL-AF9 miR-E PCGF1 engraftment in BM, SP, LV and analysis of GFP induction. (A) FACS analysis of bone marrow (BM), spleen (SP) and liver (LV) of primary CB MLL-AF9 (mBIueberry) miR-E PCGF1 (mCherry) mouse. (B) FACS plots of engraftment levels in BM of MLL-AF9 mBlueberry/miR-E PCGF1 mCherry mice are illustrated and GFP levels were divided into negative (neg), mid and bright populations. Mouse #1 was administered doxycycline for 5 days before sacrifice to determine max GFP induction in the BM. (C) Example of chimerism level in BM (no doxycycline #1, 5 days dox), gated for top, middle and bottom mCherry+ population and further analyzed for GFP percentage.

